# Altered virome structrue and function characterization in *Helicobacter pylori*-driven colorectal carcinogenesis and *H. pylori* eradication

**DOI:** 10.1101/2022.07.03.498559

**Authors:** Shiqi Luo, Jinling Xue, Jinlong Ru, Mohammadali Khan Mirzaei, Anna Ralser, Raquel Mejías-Luque, Markus Gerhard, Li Deng

**Affiliations:** Institute of Virology, Helmholtz Centre Munich – German Research Center for Environmental Health, 85764 Neuherberg, Germany; Chair of Prevention of Microbial Diseases, School of Life Sciences Weihenstephan, Technical University of Munich, Freising, Germany; Institute for Medical Microbiology, Immunology and Hygiene, Technical University of Munich, Munich, 81675, Germany; German Center for Infection Research (DZIF), Munich Partner Site, Munich, 81675, Germany

## Abstract

The understanding of gut virome and its role in *Helicobacter pylori*-driven colorectal cancer (CRC), as well as the long-term impact of *H. pylori* eradication via antibiotic treatment on it could contribute to better understanding the mechanisms of the disruption of gut bacteriome homeostasis involved in *H. pylori*-driven colorectal carcinogenesis and antibiotic therapy for *H. pylori* eradication. In the dynamic analysis of viral genome shotgun metagenomic of samples from lower gastrointestinal tract of the *Apc^+/1638N^* and C57BL/6 mice with *H. pylori* infection and eradication, stable viral abundance and replacement of bursted unique viral contigs in infected and uninfected *Apc^+/1638N^* mice were observed. Temperate phages, which encoding comprehensive microbial functional genes and targeting various susceptible hosts, were expanded extremely prior to cancer exacerbation. In addition, short-term antibiotic exposure for *H. pylori* eradication was able to alter the gut virome and thrive the antibiotic resistance genes (ARGs) in the viral genome for at least 6 months. Collectively, these results point toward a potential role of the altered, but dynamically balanced gut virome, characterized by the expanded temperate phages, in contributing to the *H. pylori*-driven CRC, and indicate that viral genome may act as ARG reservoir for the antibiotic resistance of bacteria after the antibiotics therapy to *H. pylori* eradication.

## Introduction

*H. pylori* infection is extremely common and affects more than 50% of the world population, as the cause of several gastric and extragastric diseases (Gravina et al., 2018). The epidemiological findings and meta-analysis have implied a close correlation between CRC and *H. pylori* infection (Kim et al., 2017; Butt and Epplein, 2019; Zuo et al., 2020). To better understand the mechanisms of causal effect of *H. pylori* on CRC, Ralser et al. recently conducted a series of analyses of *H. pylori*-induced changes in gut environment on *Apc*-mutant mouse models and C57BL/6 mice, and further verified the findings in human samples (Ralser et al., 2022). *H. pylori* infection was found to cause the deregulation of enteric immunity, and significantly alter the bacterial community structures, characterized by the mucus-degrading microbiota expansion (Ralser et al., 2022). In short, the homeostasis in the gut would be disrupted by the *H. pylori* invasion and contribute to colorectal carcinogenesis.

Beyond the bacteria, virus is also a major component of the gut microbiota, involved in the maintenance of the gut homeostasis. Phages are viruses infecting bacteria, making up the vast majority (97.7%) of the gut viral genome (Gregory et al., 2020). Most of them can be classified as virulent or temperate phages based on their life cycles, Upon infection, the virulent phages take over the machinery of the host cell to produce and finally lyse the host cell, whilst the temperate phages have not only lytic but also lysogenic cycles, in which phages incorporate their genomes into host chromosomes and proliferate with them rather than lysing (Howard-Varona et al., 2017). Therefore, temperate phages are regarded as natural vectors for gene transmission among susceptible bacteria and play important roles in bacterial pathogenicity (Boyd, 2012; Cuenca et al., 2016). Additionally, several studies demonstrate that phages could kill or inhibit non-target, bystander and competitor bacteria by cascading effects (Hsu et al., 2019), type VII secretion system with sub-lethal antibiotic exposure (Chatterjee et al., 2021), type V and VI secretion system with producing polymorphic toxins (Jamet et al., 2017), respectively. Together, the impact of phage population changes on the bacterial community is sophisticated and their interactions could alter the pathogenesis.

Accumulating researches observed the changes in the fecal virome, especially the phage population, in the CRC patients (Hannigan et al., 2018; Nakatsu et al., 2018; Handley and Devkota, 2019; Marônek et al., 2020). It has been suggested that phages may play a causative role in CRC by altering the bacterial populations of the intestine such that pathogenic bacteria can thrive and form biofilms (Flynn et al., 2016; Hannigan et al., 2018). And the pathological activation of some phages is assumed as a vital trigger (Hannigan et al., 2018). Meanwhile, several phages, including *Streptococcus* phage SpSL1, *Enterobacteria* phage HK544, Punalikevirus, Lambdalikevirus and Mulikevirus were found to increase in patients with CRC (Nakatsu et al., 2018; Emlet et al., 2020). However, no previous study has investigated the alteration of gut virome and its potential function on carcinogenesis accelerated by the invasion of *H. pylori*.

Furthermore, considering the known risk of *H. pylori* infection for multi-diseases, including CRC, early diagnostics and eradication of it are frequently prescribed as useful disease prevention (Malfertheiner et al., 2017; Liou et al., 2020; Ding et al., 2022). The currently recommended eradication regimens still rely on antibiotics (Suzuki et al., 2022). However, the eradication rate of *H. pylori* is declining globally every year, mainly due to the rising prevalence of antibiotic resistance (Thung et al., 2016; Tang et al., 2020). Two clinical studies report that *H. pylori* eradication resulted in antibiotic resistance genes (ARGs) increases in the gut bacterial genome and the altered bacteriome (Guo et al., 2020; Wang et al., 2022), supporting the statements in the Maastricht V/Florence Consensus Report (Malfertheiner et al., 2017). Additionally, there are growing evidence for the potential role on the mobilization of ARGs of phages in environmental, food and human samples (Calero-Cáceres and Balcázar, 2019; Kang et al., 2021; Sala-Comorera et al., 2021; Strange et al., 2021; Anand et al.). And given that Kang et al. further proved the expansion and persistence of ARGs in human gut microbiota and the specialized transduction of ARGs in phage genome following antibiotic treatment (Kang et al., 2021), phages were thereby presumed to act as the important vector to promote the antibiotic resistance in the bacteriome. Yet, research to date has not determined the impact of *H. pylori* eradication on gut virome, not to mention on ARGs in phage genome.

To address these questions, we have conducted shotgun metagenomic sequencing on virome of cecal, stool and intestinal wash samples from both *Apc^+/1638N^* and C57BL/6 mice with and without *H. pylori* infection and eradication at different timepoints to 1. characterize the changes in viral community before and after seeable tumor development and identify the microbial function genes encoded by the phages related to carcinogenesis; 2. investigate the dynamical changes of gut virome composition and the ARGs embedding in the phage genome after *H. pylori* eradication therapy.

## Material and methods

### Study Design and Sample Collection

*Apc^+/1638N^* mice were provided by Prof. Klaus-Peter Janssen (Klinikum rechts der Isar, München) and bred under specific pathogen-free conditions in the animal facility at the Technical University of Munich. 6-week old C57BL/6 mice were purchased from Envigo RMS GmbH and acclimatized to the animal facility for 1-2 weeks. All animal experiments were conducted in compliance with European guidelines for the care and use of laboratory animals and were approved by the Bavarian Government (Regierung von Oberbayern, Az.55.2-1-54-2532-161-2017).

After the acclimation period, 6-8 weeks old *Apc^+/1638N^* and C57BL/6 mice were orally gavaged twice within 72 hours with 2 × 10^8^ *H. pylori* PMSS1. The cecum, stool, and intestinal wash samples from animals at weeks 12, and 24 after infection were subjected to downstream analysis as phase 1 study (Fig. 1A). In phase 2, the infected *Apc^+/1638N^* and C57BL/6 mice were orally gavaged twice daily for 7 consecutive days with an antibiotic cocktail containing clarithromycin (Eberth) (7.15 mg/kg/day), metronidazole (Carl Roth) (14.2 mg/kg/day) and the proton-pump inhibitor omeprazole (Carl Roth) (400 μmol/kg/day) from 4-week *pi*. The cecum, stool, and intestinal wash samples from C57BL/6 mice at 4-, 12-, and 24-week *pe* and those samples from *Apc^+/1638N^* mice at 24-week *pe* were subjected to downstream analysis (Fig. 1A). Samples were stored at −80 °C until further DNA extraction.

**Figure 1.**
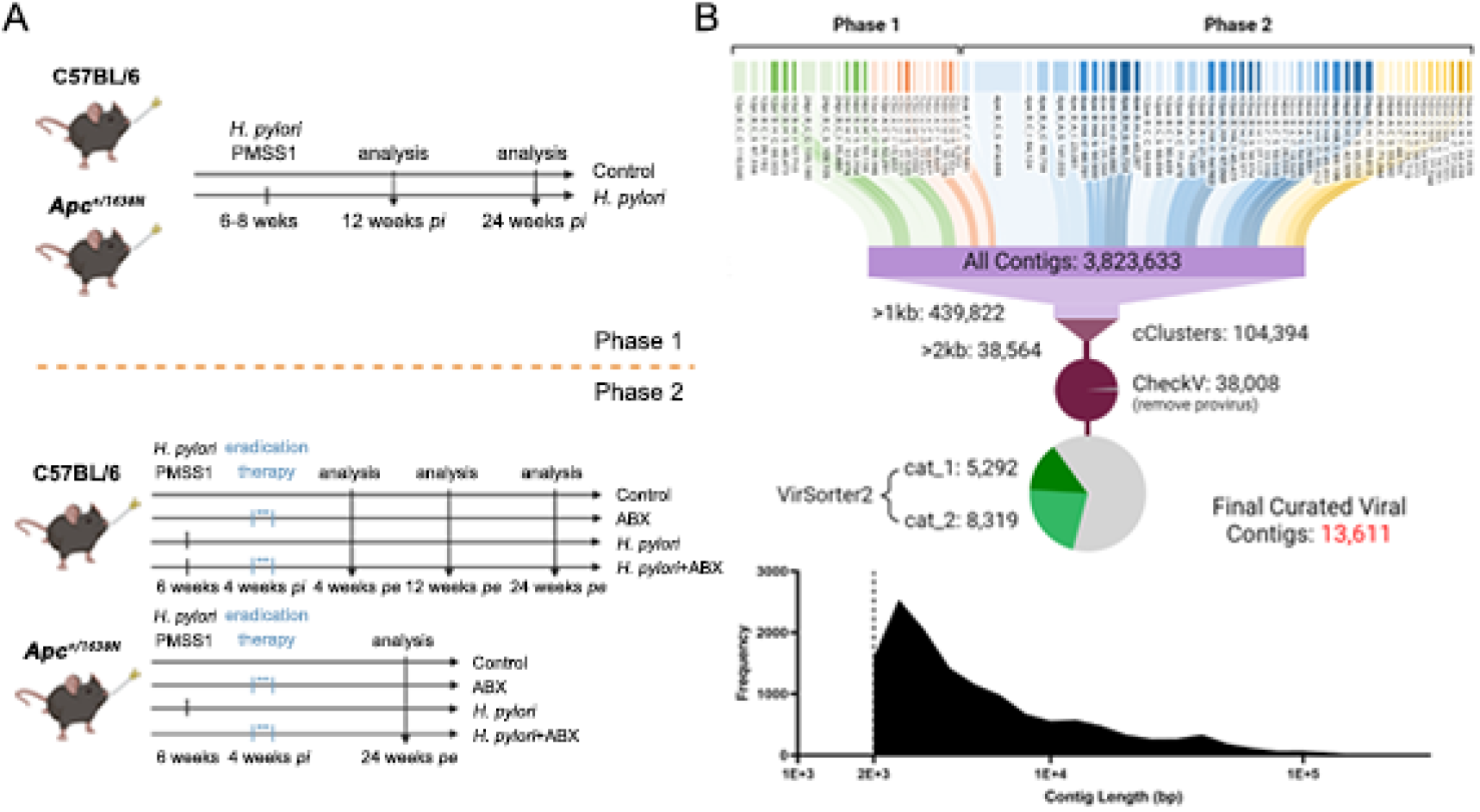
Overview of the experiment design and the viral metagenomic data. (A) Experimental setting of phases 1 and 2. (B) Generation and curation of *de novo*-assembled VCs.

### Virome Preparation and Metagenomic Sequencing

At least three samples from each group were pooled before DNA extraction to generate sufficient viral DNA for sequencing. The samples were vortexed vigorously at 4 °C overnight, then centrifugated at 4000 g, 30 min at 4 °C to obtain the supernatants. After being filtered through 0.22 μm filters (Lot No. ROCB29300, Merck Millipore), the supernatants, which were used to extract viral DNA, were concentrated to 5 mL by 10 kDa Amicon^®^ Ultra Centrifugal Filters (Lot No. R9EA18187, Merck Millipore) and then applied for CsCl density gradient centrifugation at 24,000 rpm, for 4 h at 4 °C to remove any small microbial cells. The gradient was produced using 1.2, 1.4, 1.5, and 1.65 g/cm^3^ CsCl into 14×95 mm Thinwall Ultra-clear tubes (Lot No. 344060, Beckmann). The 1.39 to 1.51 g/cm^3^ range from the gradient was collected and concentrated to less than 100 μL by 10 kDa Amicon^®^ Ultra Centrifugal Filters.

The samples were treated with Turbo DNAse (2 U/μL, Lot No. AM2238, Invitrogen) treatment for 1 h at 37 °C and followed by treatment with lysis buffer (700 μL KOH stock (0.43 g/10 mL), 430 μL DDT stock (0.8 g/10 mL), and 370 μL H2O, pH = 12) at room temperature for 10 min, −80 °C for 2 h and 55 °C for 5 min. Lysed viral-like particles were then treated for 30 min at 55 °C with Proteinase K (20 mg/mL, Lot No. 1112907, Invitrogen) to digest the viral protein capsid. To obtain viral DNA with high purity, AMPure XP beads (Lot No. A63882, Beckman Coulter) were subsequently added to samples and incubated at room temperature for 15 min and followed by twice washes with 80% ethanol. DNA was then eluted from beads with 35 μL Tris buffer (10 mM, pH = 9.8). The concentration and quality were determined using the Qubit dsDNA HS kit (Lot No. Q32854, Invitrogen) and Agilent 5400 fragment analyzer system (Agilent Technologies). The library was prepared using Nextera XT DNA Library Preparation kit (Lot No. FC-131-1096, Illumina) and sequencing was performed on an Illumina Novoseq 6000 platform.

### Bioinformatic Analysis

Illumina sequencing reads were downloaded and processed as follows. Raw reads were filtered with fastp (v0.23.1) (Chen et al., 2018) to remove adaptors and low-quality bases. Host contaminations were removed by mapping clean reads to a masked host reference genome (Handley, Scott A, 2020) using bbmap.sh script (Bushnell, 2015) Briefly, host reference genome was downloaded from NCBI genome database. Viral sequences were downloaded from viral RefSeq and neighbor nucleotide records in NCBI nucleotide database, and shred using shred.sh script (Bushnell, 2015). The shredded virus sequences were then mapped to the host reference genome using bbmap.sh script. Mapped regions were masked using bbmask.sh script (Bushnell, 2015). Unmapped reads were extracted for downstream analysis.

Duplicate reads were removed using dedupe.sh script (v38.90) (Bushnell, 2015). Deduplicated reads were assembled into contigs using metaSPAdes (v3.15.4) (Nurk et al., 2017) with default parameters except that k-mer size were set to 21, 33, 55, 77, 99. Contigs from all samples were merged to a single contig library file (cclib). The size of cclib were reduced by removing those shorter than 1 kbp and 100% similarity duplicates using seqkit (Shen et al., 2016) and dedupe.sh script. Remaining contigs were processed using CheckV (v0.8.1) (Nayfach et al., 2021) with in-house script to extract viral region from contigs that contain provirus. To further reduce the redundancy of the contig library, we clustered contigs following the “rapid genome clustering based on pairwise ANI” protocol in CheckV. Contigs were clustered if they shared higher than 95% identity and higher than 80% coverage. The longest contigs in each cluster were selected as a representative contig and formed the non-redundant contig library (nrclib). Contigs in nrclib were sent to VirSorter2 (v2.2.3) (Guo et al., 2021) for viral detection. Contigs classified into category 1 and 2 by VirSorter2 were included in this study as viral contigs (VCs). For each VC, ORFs were predicted using Prodigal (v2.6.3) (Hyatt et al., 2012) and provided to vConTACT2 (v0.11.1) (Bin Jang et al., 2019), CAT (5.2.3) (von Meijenfeldt et al., 2019) and Demovir script (https://github.com/feargalr/Demovir) for taxonomy annotation.

Clean reads were mapped to nrclib using minimap2. Coverage and depth of contigs were calculated using CoverM (v0.6.1) (https://github.com/wwood/CoverM). Transcripts per million (TPM) for each contig was calculated and normalized in each sample to obtain the relative abundance of each contig. VCs were fed into bacplip (v0.9.6) (Hockenberry and Wilke, 2021), Deephage (v1.0) (Wu et al., 2021) and an in-house software, Replidec (Peng, unpublished) for phage life cycle determination and the final vote was given by at least two tools. Auxiliary metabolic genes (AMGs) embedding in VCs were annotated using DRAM-v (Shaffer et al., 2020). The VCs were applied to CARD database (Alcock et al., 2020) for annotating the antibiotic resistance gene homolog (ARG) and antibiotic-specific resistance gene homolog (AsRG), which were ARG homologs with proven resistance to each antibiotic given to the mice. These AsRG were verified by CARD (Alcock et al., 2020) and only genes listed as “confers_resistance_to_drug” in “Sub-Term(s)” of each antibiotic were kept. The phage-bacterial host interactions were predicted by VirHostMatcher-Net (Wang et al., 2020). Bacterial hosts were predicted for VCs with score higher than 90% according to VirHostMatcher-Net. VCs, which were able to infect more than one host, were classified as polyvalent VCs.

### Statistics Analysis

Alpha (Shannon and Chao1 index) and beta diversities were calculated at VC level in vegan package in R (v4.1.2). Significant differences in mice with different treatments identified via constrained ordination were confirmed using PERMANOVA (permutational multivariate analysis of variance) via the *adonis* function in vegan package. The enriched and depleted VCs in the samples from *Apc^+/1638N^* infected by *H. pylori* PMSS1 were identified by using DESeq2 package in R. All plots were plotted with ggplot2 package.

## Results

### *H. pylori*-driven CRC is characterized by an altered, but dynamically balanced viral community

The fact that *H. pylori* infection accelerates colorectal carcinogenesis, and early antibiotic eradication of *H. pylori* infection reduces the tumor incidence to the level of uninfected controls have been demonstrated before (Ralser et al., 2022).

After sequencing, 13,611 VCs with the length of 2-5 Kb (60%), 5-10 kb (18.2%), 10-50 kb (19.1%), 50-268 kb (2.7%), were assembled from the cecum, stool and intestinal wash samples of animals collected as the setting of phase 1 and 2 in Fig. 1A, and employed in the downstream analysis (Fig. 1B).

The virome was naturally different in *Apc^+/1638N^* and C57BL/6 mice as less virome diversity and richness were seen in *Apc^+/1638N^* mice compared to in C57BL/6 mice regardless of infection (Fig. 2A). *H. pylori* infection only reduced the virome diversity and richness in C57BL/6 mice (Fig. 2A). However, the divergence in the virome of *Apc^+/1638N^* mice induced by infection was significantly greater than in the virome of C57BL/6 mice (Fig. 2B). Considering that 12.82% and 23.4% of the 10031 VCs existed uniquely and respectively in infected and uninfected *Apc^+/1638N^* mice, and they were only 2.69% and 6.19% of 13107 VCs in C57BL/6 mice (Fig. 2C), the divergence in virome could be attributed to the replacements of many unique VCs in *Apc^+/1638N^* mice with *H. pylori* infection. Together with the observation of tumors in *Apc^+/1638N^* mice solely, it has demonstrated that the *H. pylori-* driven CRC is associated with a dynamically balanced viral community, rather than a viral community with straightforward reductions in diversity and richness.

**Figure 2.**
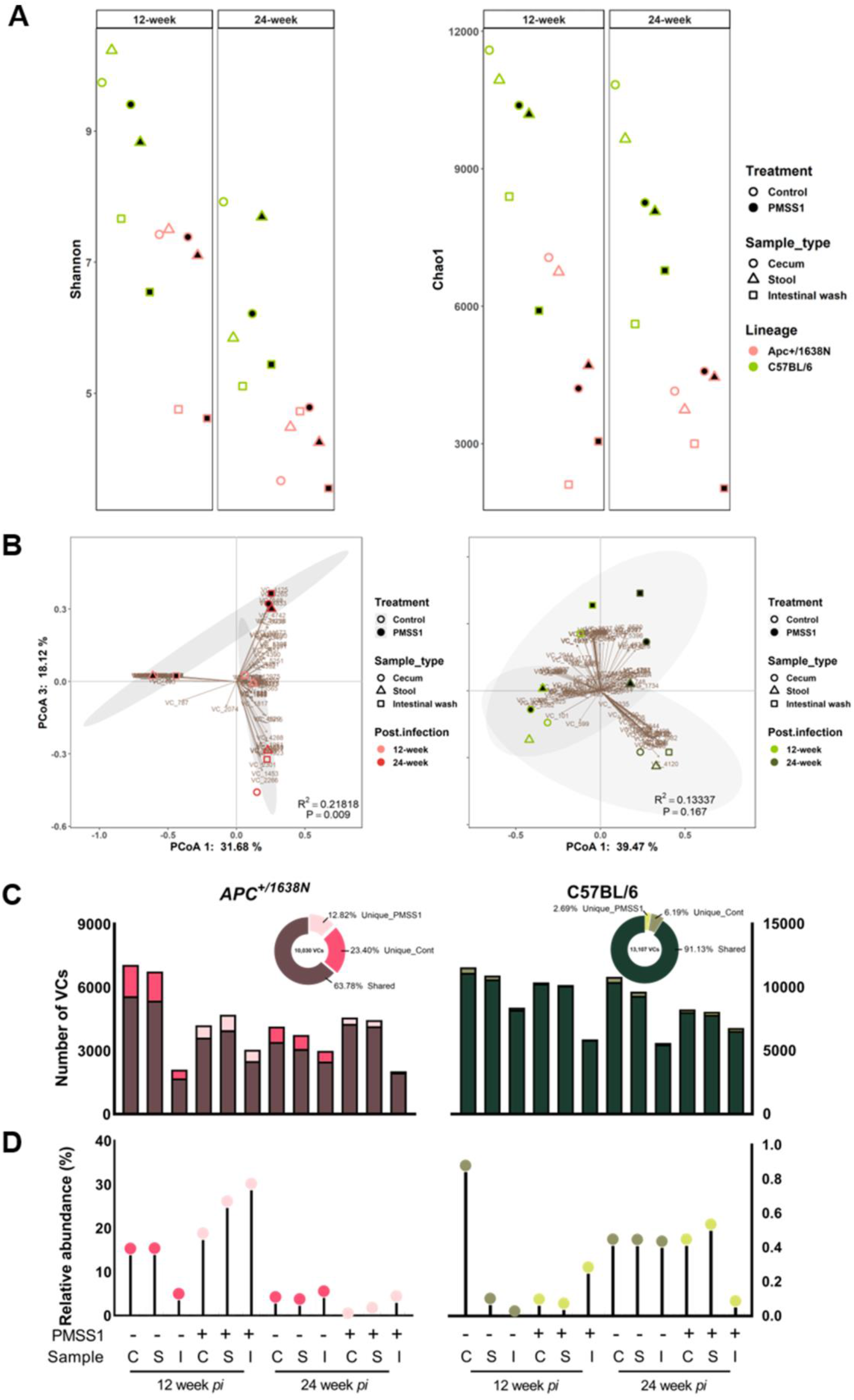
Structure of viral community of mice from phase 1. (A) α-diversity and (B) PCoA of virome in the cecum, stool and intestinal wash of *Apc^+/1638N^* and C57BL/6 mice with and without *H. pylori* infection at 12-, 24-week *pi*. (C-D) Distribution of shared and unique VCs: 10031, 13107 detected VCs in *Apc^+/1638N^* and C57BL/6 mice were assigned into 3 categories based on overlapping characteristics among samples with and without PMSS1 infection. 1286 VCs were detected only in PMSS1 treated *Apc^+/1638N^* mice and 2347 VCs were detected only in non-treated *Apc^+/1638N^* mice. There were 352 and 811 VCs detected in PMSS1 treated and non-treated C57BL/6 mice. (C) The absolute number of detected VCs in relationship to sharing characteristics. (D) The relative abundance of unique VCs in each sample. One dot/bar represents at least three mice.

### Expanding Temperate Phages in *Apc^+/1638N^* Gut Virome May Promote the Exacerbation of *H. pylori*-Driven CRC via Enhancing Bacterial Pathogenicity

The ratio of temperate and virulent phage in virome was computed based on the final vote by tools using three different algorithms (Fig. 3A). It showed that the absolute domination of temperate phages in gut (79.2% in cecal, 74.4% in stool and 91.4% in intestinal wash) virome occurred in *Apc^+/1638N^* mice at 12-week *pi*, the early stage of *H. pylori*-driven CRC, but not in other mice or timepoints (Fig. 3A). We next compare the relative abundance of VCs across the samples from *Apc^+/1638N^* mice to screen the expanded temperate VCs involved in the exacerbation of *H. pylori*-driven CRC at 12-week *pi*, namely, the insertion of the comparisons of increased temperate phages in infected and uninfected mice at 12-week *pi*, and in infected mice at 12- and 24-week *pi* (Fig. 3B). After clustering the main microbial functional genes embedding in the enriched temperate VCs and virulent VCs in the comparisons of infected and uninfected mice at 12-week *pi*, infected and uninfected mice at 24-week *pi*, and infected mice at 12- and 24-week *pi*, 124 genes were found to dominate the main functional role of the expanded temperate VCs involved in the exacerbation of *H. pylori*-driven CRC at 12-week *pi*, and completely different from the dominated functional genes encoded by the temperate VCs enriched in infected mice at 24-week *pi* (Fig. 3C, details in Fig. S1). Together with the stable performance of the comprehensive function behind these genes among all sample types, the temperate VCs harboring these genes are suggested to be involved in triggering the aggravation of *H. pylori*-driven CRC by manipulating the comprehensive bacterial bioactivities, including genetic information processing, signaling and cellular processes, metabolism and so on (Fig. 3D).

**Figure 3.**
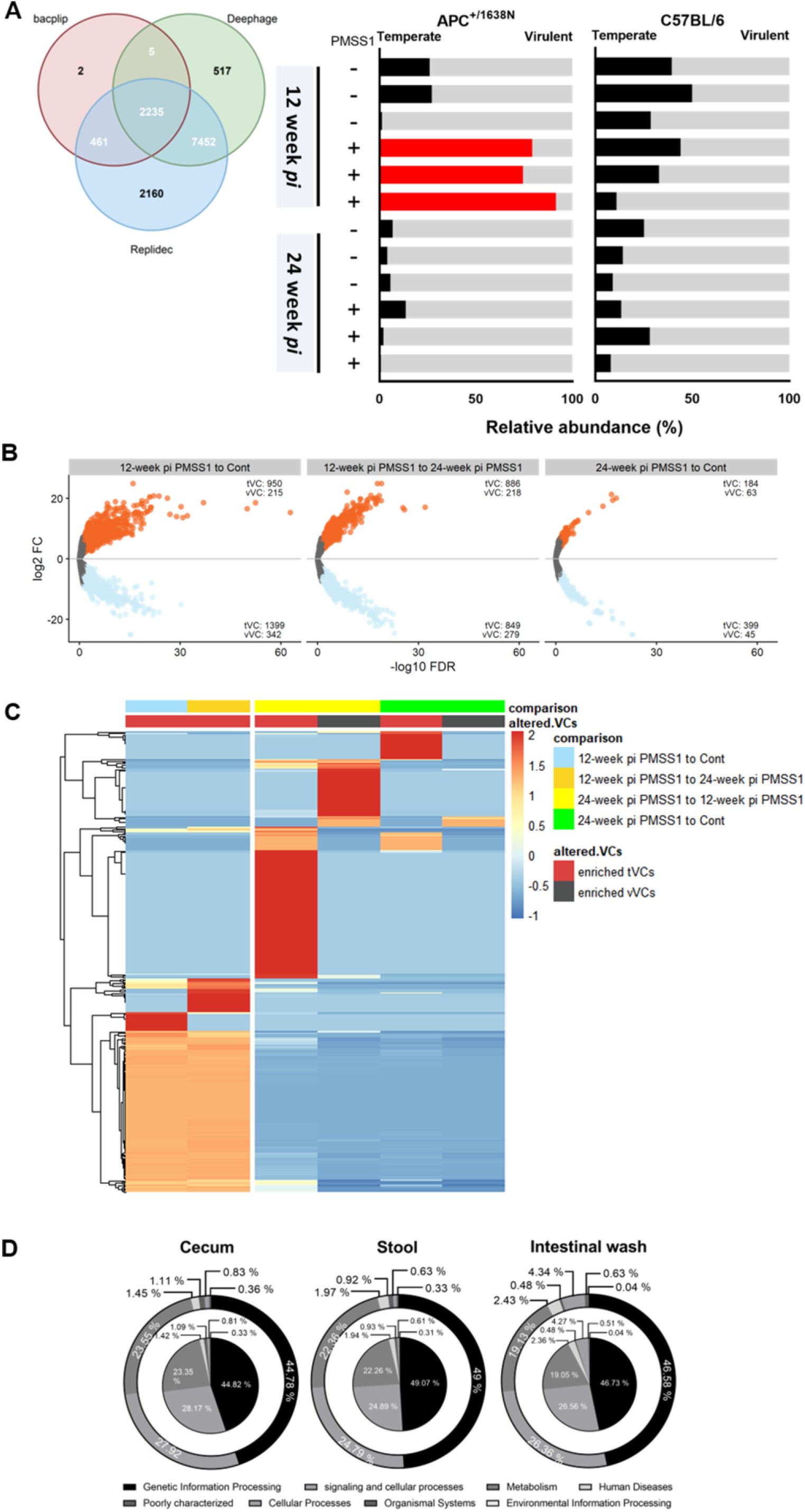
Expanded temperate phages in *Apc^+/1638N^* mice at 12-week *pi* and their microbial functional genes. (A) Ratio of temperate and virulent phage in gut virome of *Apc^+/1638N^* mice. (B) The enriched and depleted, temperate and virulent viral contigs in cecum, stool and intestinal wash of *H. pylori* treated *Apc^+/1638N^* mice compared to non-treated mice at 12-, 24-week *pi*, and of *H. pylori* treated *Apc^+/1638N^* mice at 12-week *pi* compared to 24-week *pi*. (C) The function annotation of the enriched temperate viral contigs (tVCs) and virulent viral contigs (vVCs) in the six groups of comparisons. (D) The function distribution of the intersection of functional genes embedded in the enriched tVCs in the comparisons of infected and uninfected mice at 12-week *pi*, and infected mice at 12- and 24-week *pi*, representing the main microbial functions carried by the tVCs remarkably enriched in cecum, stool and intestinal wash of *Apc^+/1638N^* mice in the early stage of CRC development. Genes are annotated in KEGG (first level). Inner pies represent the data for infected and uninfected mice at 12-week *pi* whilst outer pies for infected mice at 12- and 24-week *pi*.

In addition, evidence showed that these enriched VCs tended to manipulate the bacterial species belonging to the bacterial genera, which have been reported to have strong associations with CRC development, such as *Clostridium, Enterococcus, Staphylococcus* and *Clostridioides* (Fig. 4A). While being infected, these pathogenic bacteria might be supplemented with the various functional genes shown in Fig. S1, including the virulence genes (e.g. *flgJ* (Coloma-Rivero et al., 2020)*, MazF* (Schifano et al., 2016)) and controller of virulence genes transfer (e.g. *dut* (Tormo-Más et al., 2013)). Interestingly, the well-known residential probiotics from genus *Lactobacillus*, as well as butyrate-producing bacterium from *Blautia* and *Butyrivibrio* are the potential hosts of the expanded temperate VCs involved in the exacerbation of *H. pylori*-driven CRC (Fig. 4A), implying that the known loss of these probiotics in CRC development (Chen et al., 2012; Borges-Canha et al., 2015; Drewes et al., 2016) could be attributed to these enriched temperate VCs at the early stage. Moreover, these enriched temperate VCs showed higher polyvalence (14.1% in the enriched temperate phages in 12-week post infected animals compared to uninfected animals; 12.6% in the enriched temperate phages in 12-week post infected animals compared to 24-week post infected animals) at the early stage of *H. pylori*-driven CRC formation (Fig. 4B), indicating their high tendency to transfer the bacterial functional genes across the susceptible hosts to exacerbate the carcinogenesis.

**Figure 4.**
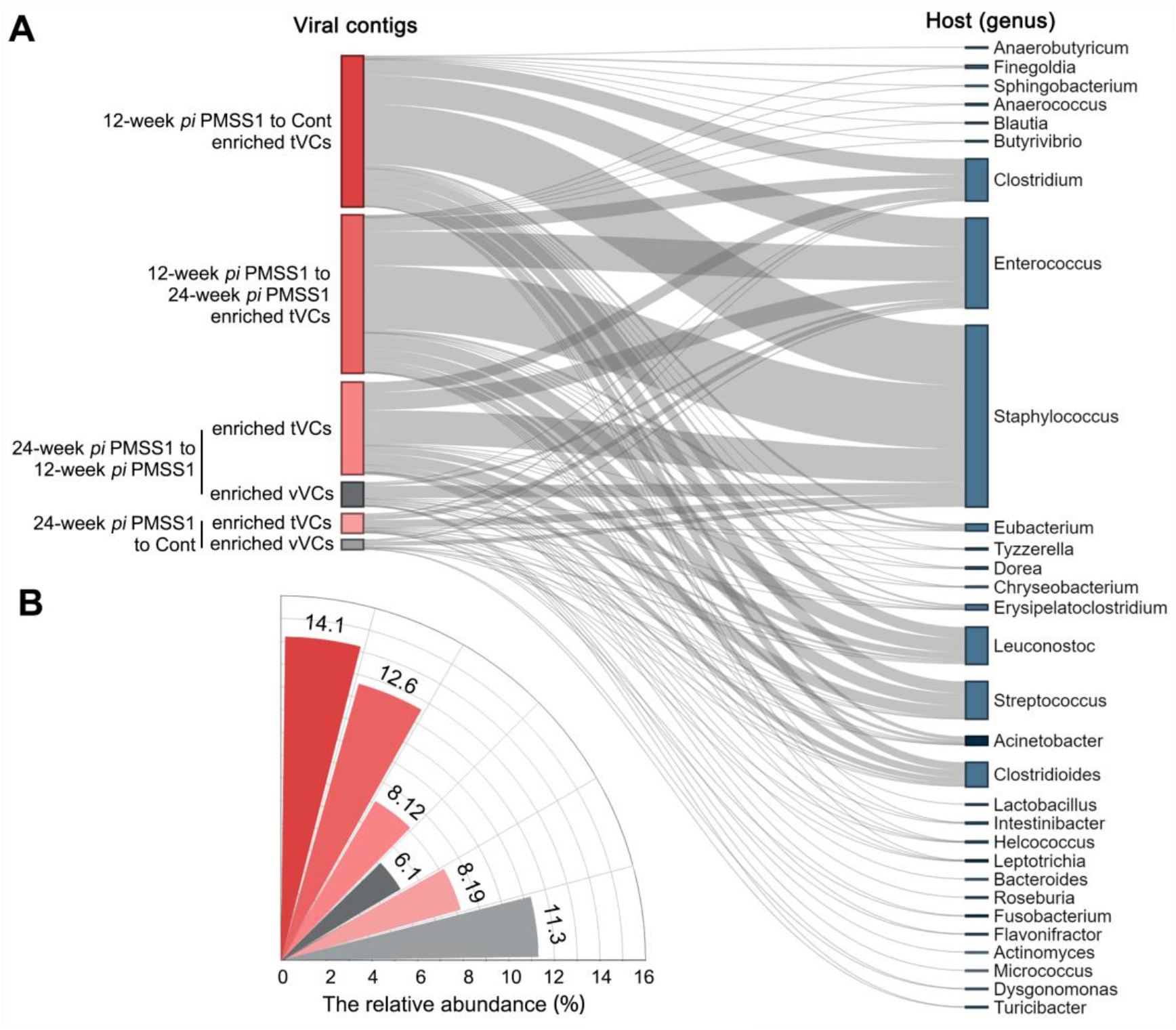
Predicted bacterial hosts of the enriched phages *Apc^+/1638N^* mice. (A) Predicted virus-host linkages of the enriched tVCs and vVCs in *Apc^+/1638N^* mice. (B) The relative abundance of polyvalent VCs within the six VC subsets, respectively. Polyvalence was found to be higher in enriched tVCs in mice at 12-week *pi* with *H. pylori*.

### Altered Virome Structure and Expanded Antibiotic Resistance Genes in Viral Genome after Antibiotics Treatment of *H. pylori*

Antibiotic treatment exacerbated the reduction of virome diversity and richness triggered by *H. pylori* infection in C57BL/6 mice at 4-week *pe*. Similar reduction of virome diversity brought by *H. pylori* plus antibiotic treatment was found in *Apc^+/1638N^* mice at 24-week *pe* (Fig. 5A). In addition, the divergent viral communities were not only observed in *Apc^+/1638N^* mice (Fig. 2B), but also C57BL/6 mice. This divergence of gut virome in C57BL/6 mice was primarily contributed by antibiotics treatment because samples from the mice without treatment and only *H. pylori* treatment remained tightly grouped (Fig. 5B), similar with the previous finding (Fig. 2B), whilst it was caused by *H. pylori* infection and the subsequent antibiotic treatment in *Apc^+/1638N^* mice. These findings showed that antibiotic therapy after *H. pylori* infection not only eradicated *H. pylori*, but also altered the gut virome in both *Apc^+/1638N^* and C57BL/6 mice for at least 24 weeks. The alteration is irreversible without further intervention.

**Figure 5.**
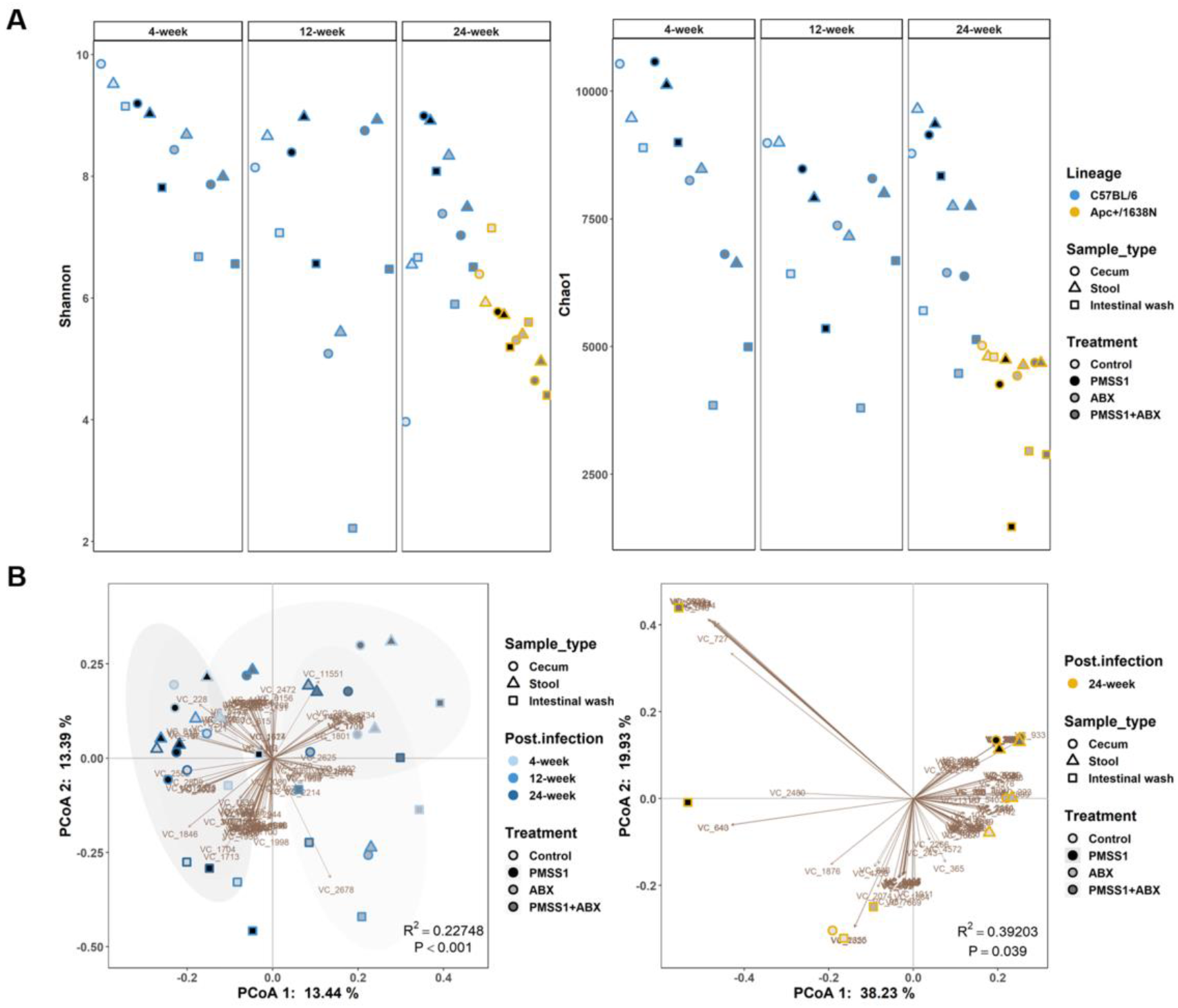
Structure of viral community of mice from phase 2. (A) α-diversity and (B) PCoA of virome in cecum, stool and intestinal wash of *Apc^+/1638N^* and C57BL/6 mice with presence and absence of antibiotics (clarithromycin, metronidazole and omeprazole) treatment and *H. pylori* infection at 4, 12, 24-week post eradication. Each dot represents at least three mice.

For further investigation of the implications of the virome alteration on the gut bacteriome, we first looked at whether the antibiotic resistance genes carried by the VCs increased after the therapy. The normalized number of temperate VCs harboring antibiotic resistance gene homologous (ARGs) was slightly increased in antibiotic-treated C57BL/6 mice after 4- and 12-week *pe* (Table S1) and the relative abundance of temperate VCs harboring ARGs in the mice with antibiotic therapy after *H. pylori* infection were higher (19.60% in cecum, 12.57% in stool, 5.44% in intestinal wash at 4-week *pe*; 18.70% in cecum, 18.98% in stool, 3.59% in intestinal wash at 12-week *pe*) than those mice with *H. pylori* infection only (7.43% in cecum, 8.55% in stool, 3.11% in intestinal wash at 4-week *pe*; 14.29% in cecum, 17.80% in stool, 3.93% in intestinal wash at 12-week *pe*) or antibiotics therapy only (4.25% in cecum, 6.09% in stool, 1.20% in intestinal wash at 4-week *pe*; 7.30% in cecum, 6.70% in stool, 3.67% in intestinal wash at 12-week *pe*), and the abundance gap continued to diminish till the 24-week *pe*. in C57BL/6 mice (Table S2). However, the higher relative abundance of temperate VCs harboring ARGs in cecum and stool of *Apc^+/1638N^* mice administrated with *H. pylori* plus antibiotics compared to with only *H. pylori* or antibiotics (7.22% vs. 1.46% and 2.51% in cecum, 8.72% vs. 1.96% and 2.93% in stool) still existed at 24-week *pe* (Table S2). In addition, there were at least 17 ARGs expanding in viral genome in the gut of the *H. pylori*-eradicated *Apc^+/1638N^* mice (Fig. 6A). Among them, we found the antibiotic-specific resistance gene homologous (AsRGs) - *Erm(k), Erm(30), Erm(34)* and *msbA*, which were verified to be associated with the clarithromycin, metronidazole-resistance, respectively. The total relative abundance of temperate VCs harboring these four AsRGs were higher in the C57BL/6 mice with *H. pylori* plus antibiotics than with only *H. pylori* or antibiotics administration at 4-week *pe* (0.32% vs. 0.05% and 0.06% in cecum, 0.38% vs. 0.04% and 0.06% in stool, 0.07% vs. 0.02% and 0.07% in intestinal wash) and 24-week *pe* (0.20% vs. 0.06% and 0.06% in cecum, 0.11% vs. 0.09% and 0.15% in stool, 0.45% vs. 0.17% and 0.15% in intestinal wash) (Table S3). Noteworthy, the expansion of each AsRG in viral genome was different (Fig. 6A and Fig. 6B). For example, *Erm(30)* only expanded in C57BL/6 mice at 4-week *pe*, whilst *msbA* expanded at 12- and 24-week *pe*, and even longer beyond the 24 weeks (Fig. 6B). It roughly illustrated that the strength and duration of the influence caused by the antibiotic eradication of *H. pylori* on virome depended on the antibiotics utilized.

**Figure 6.**
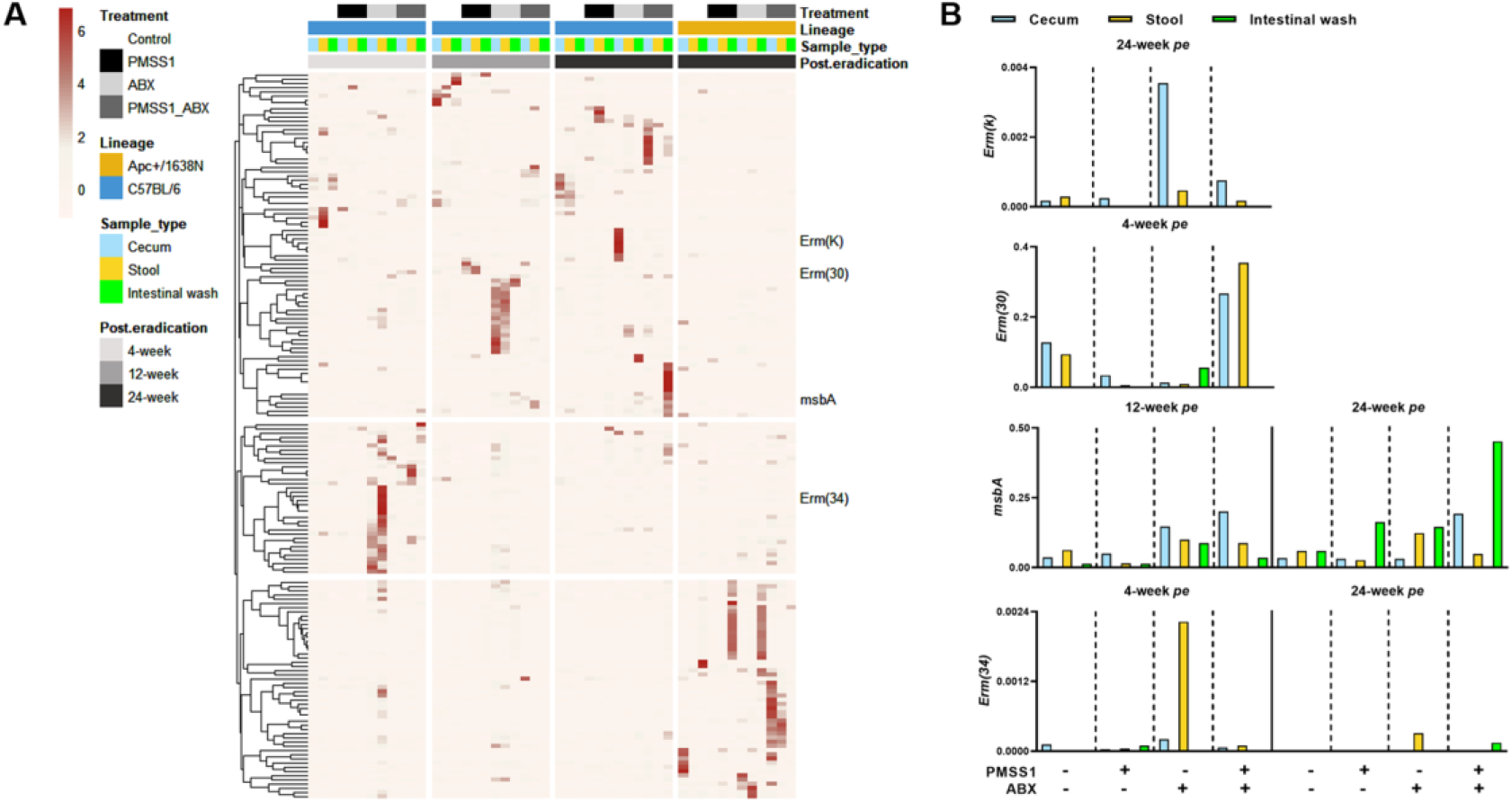
ARGs and AsRGs profile in the gut virome. (A) Heatmap of the relative abundance of temperate VCs harboring ARGs in the virome of *Apc^+/1638N^* and C57BL/6 mice with presence and absence of antibiotics treatment and *H. pylori* infection at 4-, 12-, 24-week post eradication. AsRGs, which were verified to be associated with the clarithromycin, metronidazole-resistance, were labeled in the plot. (B) The percentage of relative abundance of temperate VCs carrying AsRGs (*Erm(k), Erm(30), Erm(34) and msbA*, in the virome of C57BL/6 mice with presence and absence of antibiotics treatment and H. pylori infection at a certain time point. Each bar represents at least three mice.

Together, antibiotic eradication of *H. pylori* would change the viral community and further exert detrimental implications on the bacterial community via the antibiotic resistance genes expansion in the viral community.

## Discussion

There is mounting evidence showing that disturbances to the gut virome, especially phage populations, could lead to diseases including CRC because of phages’ capacity for mutagenesis and modulating their host functionality (Handley and Devkota, 2019; Broecker and Moelling, 2021; Li et al., 2022; Tiamani et al., 2022). Here we show that, in the model of *H. pylori*-driven CRC, the residential gut viral members were replaced by distinct phages. Interestingly, the temperate phages encoding a variety of microbial functional genes dominated the gut virome at the early stage of *H. pylori*-driven CRC. Our findings support a working hypothesis for carcinogenesis by the phage-modulated bacterial community.

The alterations of fecal virome have been emphasized to be linked with CRC, but the tendencies are not consistent in the previous researches (Hannigan et al., 2018; Nakatsu et al., 2018; Li et al., 2022), and this study (Fig. 2 and Fig. 3). It was mainly because the initial trigger of CRC in the previous animal study was the chemical compound - azoxymethane, and those in clinical researches were unknown, and likely to differ from this study, in which *H. pylori* was the certain, initial trigger. In particular, basic diversity metrics of alpha diversity (richness and Shannon diversity) and beta diversity (Bray-Curtis dissimilarity) of virome presented different tendencies in experimental models and clinical patients with CRC relative to those without CRC (Hannigan et al., 2018; Nakatsu et al., 2018; Li et al., 2022) (Fig. 2A and Fig. 2B), which implies that the difference in alpha and beta diversity of virome is insufficient for identifying healthy or cancerous states without taking into account the initial triggers, and further implies that the alteration of virome is not the uniform outcome of CRC, but more likely to involve in the pathogenesis after being activated by the initial triggers, such as *H. pylori* infection.

Besides, it is noted that *Apc-*mutant mice had significantly more unique VCs than C57BL/6 mice, particularly at 12-week *pi* (Fig. 2D), which was prior to CRC aggravation since the animals had not developed visible tumors. Interestingly, the gut virome of infected *Apc-* mutant mice was dominated by temperate phages at this period (Fig. 3A). The shift of core gut virome from virulent phages to temperate phages, which were induced from bacteria, has been reported to be indicative of IBD (Clooney et al., 2019). The expanded temperate phages preceded CRC aggravation supports the assumption of the vital role of pathological activation of some phages in CRC carcinogenesis (Hannigan et al., 2018). The pathological activation of the temperate phages is to induce the silent prophages in lysogenic state into lytic state from the bacterial host (Du Toit, 2018). In addition to the well-known antibiotic treatment (e.g., mitomycin C), the induction process could also be initiated by many kinds of stresses, including bacterial toxin and metabolite (Jancheva and Böttcher, 2021; Silpe et al., 2022), signals of quorum sensing of bacteria (Liang et al., 2020), and stress from competing bacteria (Roossinck, 2011). Specifically, under the stress from competing bacteria, the host bacteria would automatically sacrifice to release the prophages to kill the competing bacteria for the benefit of the remaining lysogenic population of bacteria (Bossi et al., 2003). The previous study has showed the competitive stress would be brought by *H. pylori* infection on probiotics, such as *Lactobacillus* spp. (Coconnier et al., 1998). Together, the observation of expansion of temperate phages prior to CRC aggravation might be attributed to the sacrifices of the residential probiotics to fight against *H. pylori* and this was reflected by the fact that several probiotics were predicted as the hosts of these expanded temperate phages (Fig. 4A).

The Polyvalence was shown to be higher in the expanded temperate phages associated with CRC aggravation (Fig. 4B). Broad host range phages can use multiple hosts to propagate and be enriched specifically in oligotrophic environments or in conditions of low host density (Yu et al., 2017), which could be triggered by *H. pylori* infection (Franceschi et al., 2014; Chen et al., 2021). Given that the hosts of these polyvalence temperate phages included CRC-associated bacteria from genus *Clostridium, Enterococcus, Staphylococcus* and *Clostridioides* (Fig. 4A) (Chattopadhyay et al., 2022), it is conceivable that these phages may act as vectors for gene transmission among bacteria and play important roles in bacterial pathogenicity in CRC aggravation.

Furthermore, to fill the gap in virome’s role in rising antibiotic resistance after *H. pylori* infection and eradication, we demonstrated the dynamic changes in ARGs and AsRGs in the viral metagenomic sequencing data. Although viromes are regarded as the ARGs reservoirs for the microbial communities (Colombo et al., 2017; Calero-Cáceres et al., 2019; Debroas and Siguret, 2019; Moon et al., 2020), a study has exposed a methodological problem that inflates ARG counts in metagenomic studies (Enault et al., 2017). To detect and discard contamination, we applied our data to the specific automated algorithm, VirSorter2, proposed by the authors exposed the methodological problem (Guo et al., 2021). The increased ARGs and AsRGs embedding in the viral genome acknowledge our assumption that phages might act as important vectors to promote antibiotic resistance in the bacteriome (Fig. 6). Consistent with that in bacteriome (Wang et al., 2022), AsRGs to different antibiotics persist in the virome for varied durations (Fig. 6B). However, our study is limited to the changes of ARGs in viral metagenomics. Further studies on microbial and bacterial metagenomics to trace the dynamical mobilization of AsRGs and functional validation of ARGs are encouraged to increase our knowledge of antibiotic treatment on *H. pylori*.

Our data and analysis first suggest that the *H. pylori*–driven CRC-associated gut viral community is altered and dynamically replaced with unique VCs; that extreme expansion of temperate phages may aggravate *H. pylori*–driven CRC by supplementing pathogenic bacterial hosts with virulent genes; and that phages mediate the resistome in animals treated with *H. pylori* and antibiotics. These findings have the potential to provide evidence in understanding how *H. pylori* infection promotes CRC, and the support of good *H. pylori* eradication, and are warranted future investigations on large-cohort humans.

## Data Availability

Sequence data have been deposited to the NCBI Sequence Read Archive (accession numbers pending).

## Contributions

S.L., L.D. and M.G. conceived and designed the study. A.R., and R.M.L. provided animal samples. J.R. assembled and sorted the raw reads of viral metagenomic data. S.L. analyzed the data and drafted the manuscript. S.L. and J.X. contributed to data interpretation. J.X., M.K.M. and L.D. contributed to revised the article. L.D. acquired the funding. All authors read and reviewed the articles.

## Acknowledgments

This work was funded by the Deutsche Forschungsgemeinschaft (DFG [German Research Foundation]) SFB1371/1-395357507 (project P09).

