## Supplementary figures and images for "Altered virome structrue and function characterization in *Helicobacter pylori*-driven colorectal carcinogenesis and *H. pylori* eradication"

### Fig. S1

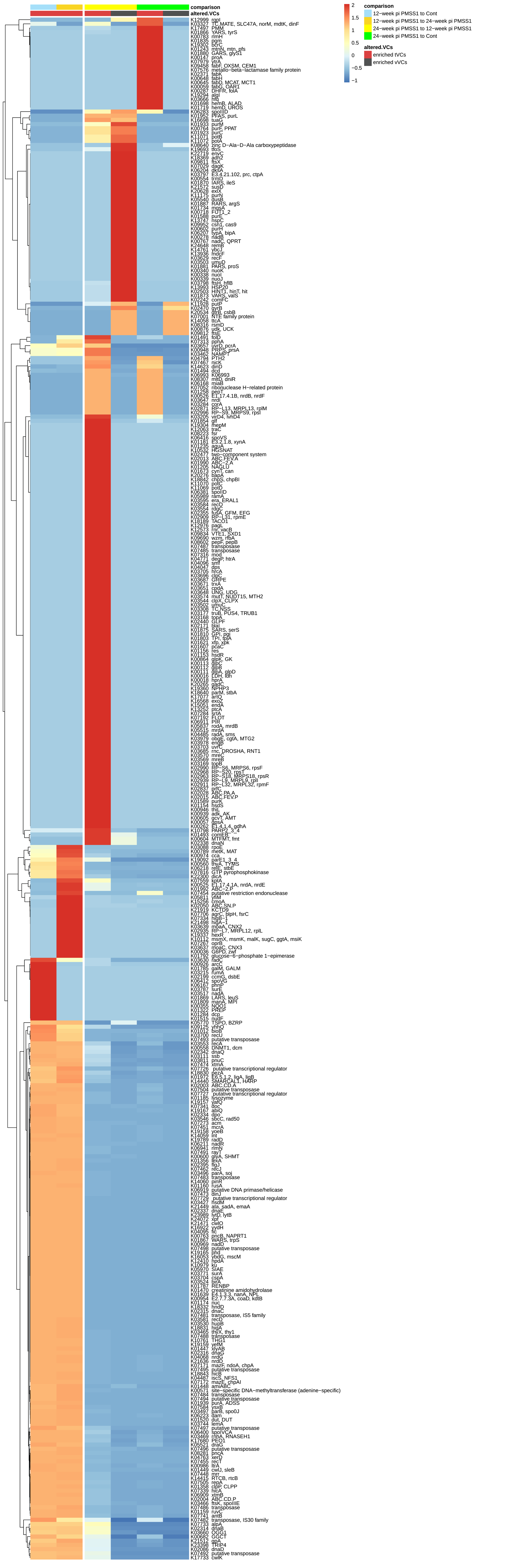
